# Evaluating the quality of the 1000 Genomes Project data

**DOI:** 10.1101/383950

**Authors:** Saurabh Belsare, Michal Sakin-Levy, Yulia Mostovoy, Steffen Durinck, Subhra Chaudhry, Ming Xiao, Andrew S. Peterson, Pui-Yan Kwok, Somasekar Seshagiri, Jeffrey D. Wall

## Abstract

Data from the 1000 Genomes project is quite often used as a reference for human genomic analysis. However, its accuracy needs to be assessed to understand the quality of predictions made using this reference. We present here an assessment of the genotype, phasing, and imputation accuracy data in the 1000 Genomes project. We compare the phased haplotype calls from the 1000 Genomes project to experimentally phased haplotypes for 28 of the same individuals sequenced using the 10X Genomics platform. We observe that phasing and imputation for rare variants are unreliable, which likely reflects the limited sample size of the 1000 Genomes project data. Further, it appears that using a population specific reference panel does not improve the accuracy of imputation over using the entire 1000 Genomes data set as a reference panel. We also note that the error rates and trends depend on the choice of definition of error, and hence any error reporting needs to take these definitions into account.

## INTRODUCTION

The 1000 Genomes Project (1KGP) was designed to provide a comprehensive description of human genetic variation through sequencing multiple individuals^1-3^. Specifically, the 1KGP provides a list of variants and haplotypes that can be used for evolutionary, functional and biomedical studies of human genetics. Over the three phases of the 1KGP, a total of 2504 individuals across 26 populations were sequenced. These populations were classified into 5 major continental groups: Africa (AFR), America (AMR), Europe (EUR), East Asia (EAS), and South Asia (SAS). The 1KGP data was generated using a combination of multiple sequencing approaches, including low coverage whole genome sequencing with mean depth of 7.4X, deep exome sequencing with a mean depth of 65.7X, and dense microarray genotyping. In addition, a subset of individuals (427) including mother-father-child trios and parent-child duos were deep sequenced using the Complete Genomics platform at a high coverage mean depth of 47X. The project involved characterization of biallelic and multiallelic SNPs, indels, and structural variants.

Given the low depth of (sequencing) coverage for most 1KGP samples, it is unclear how accurate the imputed haplotypes are, especially for rare variants. We quantify this accuracy directly by comparing imputed genotypes and haplotypes based on low-coverage whole-genome sequence data from the 1KGP with highly accurate, experimentally determined haplotypes from 28 of the same samples. Additional motivation for our study is given below.

### Phasing

It is important to understand phase information in analyzing human genomic data. Phasing involves resolving haplotypes for sites across individual whole genome sequences. The term *‘diplomics’^4^* has been coined to describe *“scientific investigations that leverage phase information in order to understand how molecular and clinical phenotypes are influenced by unique diplotypes”.* The diplotype shows effects in function and disease related phenotypes. Multiple phenomena like allele-specific expression, compound heterozygosity, inferring human demographic history, and resolving structural variants requires an understanding of the phase of available genomic data. Phased haplotypes are also required as an intermediate step for genotype imputation.

Phasing methods can be categorized into methods which use information from multiple individuals and those which rely on information from a single individual^5^. The former are primarily computational methods, while the latter are mostly experimental approaches. Some computational approaches use information from existing population genomic databases and can be used for phasing multiple individuals. These, however, may be unable to correctly phase rare and private variants, which are not represented in the reference database used. On the other hand, some methods use information from parents or closely related individuals. These have the advantage of being able to use Identical-By-Descent (IBD) information, and allow long range phasing, but require sequencing of more individuals, which adds to the cost. A few methods which use these approaches are: PHASE^6^, fastPHASE^7^, BEAGLE^8-9^, *SHAPEIT^10-11^, EAGLE^12-13^* and *IMPUTE* v2^14^.

Experimental phasing methods, on the other hand, often involve separation of entire chromosomes followed by sequencing of short segments, which can then be computationally reconstructed to generate entire haplotypes. These methods do not need information from individuals other than the one being sequenced. These methods involve genotyping being performed separately from phasing. These methods fall into two broad categories, namely dense and sparse methods^14^. Dense methods resolve haplotypes in small blocks in great detail, where all variants in a specific region are phased. However, they do not inform the phase relationship between the haplotype blocks. These involve diluting high molecular weight DNA fragments such that fragments from at most one haplotype are present in each unit. Sparse methods can resolve phase relationships across large distances, but may not inform on the phase of each variant in a chromosome. In these methods, a low number of whole chromosomes is compartmentalized such that only one of each pair of haplotypes is present in each compartment. These compartmentalizations are followed by sequencing to generate the haplotypes.

In this work, we use phased haplotypes generated using the 10X Genomics method which uses linked-read sequencing^15^. 1 nanogram of high molecular weight genomic DNA is distributed across 100,000 droplets. This DNA is barcoded and amplified using polymerase. This tagged DNA is released from the droplets and undergoes library preparation. These libraries are processed via Illumina short-read sequencing. A computational algorithm is then used to construct phased haplotypes based on the barcodes.

### Imputation

Imputation involves the prediction of genotypes not directly assayed in a sample of individuals. Experimentally sequencing genomes to a high coverage is an expensive process. Low coverage sequencing or arrays can be used as low-cost methods for sequencing. However, these methods may lead to uncertainty in estimated genotypes (low coverage sequencing) or missing genotype values for untyped sites (arrays). Imputation can be used to obtain genotype data for missing positions using reference data and known data at a subset of positions in individuals which need to be imputed. Imputation is used to boost the power of GWAS studies^16^, fine mapping a particular region of a chromosome^17^, or performing meta-analysis^18^, which involves combining reference data from multiple reference panels.

Imputation uses a reference panel of known haplotypes with alleles known at a high density of haplotyped positions. A study/inference panel genotyped at a sparse set of positions is used for sequences which need to be imputed. Performing imputation involves two basic steps:

- Phasing genotypes at genotyped positions in the study/inference panel
- Haplotypes from the inference panel which match those in the reference panel at the positions in the study panel are assumed to match in all other positions

Various imputation algorithms perform these steps sequentially and iteratively or simultaneously.

Factors affecting the quality of the phasing and imputation are (1) size of reference panel (2) density of SNPs in reference panel (3) accuracy of called genotypes in the reference panel (4) degree of relatedness between sequences in reference panel and study sequences (5) ethnicity of the study individuals in comparison with the available reference data and (6) allele frequency of the site being phased or imputed^5^.

Multiple methods have been developed for genotype imputation^19^. fastPHASE^7^, MACH^20-21^, BEAGLE^8^’^22-23^, and *IMPUTE v2^14^* are some widely used methods for imputation.

An analysis of the imputation accuracy for the HapMap project has been performed about a decade ago^24^, but no similar detailed analysis exists for assessing the phasing and imputation of the 1000 Genomes project, particularly comparing the database against experimentally phased sequences. We present here a detailed assessment of the quality of phasing and imputation for the 1000 Genomes database, particularly as a function of minor allele frequency and inter-SNP distances for biallelic SNPs.

## MATERIAL AND METHODS

### Input Data

Processed VCFs were downloaded from the 1000 Genomes website. This data is available for each chromosome separately. To obtain agreement with the experimental data, 1000 Genomes VCFs corresponding to the GRCh38 assembly were downloaded. Experimental data was sequenced using the 10X Genomics platform for 28 individuals: 5 GM, 18 HG, and 5 NA. The GM and NA individuals were originally part of the HapMap project while the HG are from the 1000 Genomes project. Thirteen of these individuals were processed at UCSF and sequenced at Novogene, while the remaining individuals were processed and sequenced at Genentech. The populations from which each of the individuals come (as listed in the Coriell Catalog) are:

- South Asia (SAS):

- Gujarati Indians in Houston, Texas, USA (HapMap) [GIH] - GM21125*, NA20900, NA20902
- Punjabi in Lahore, Pakistan [PJL] - HG03491, HG03619
- Sri Lankan Tamil in the UK [STU] - HG03679, HG03752, HG03838*
- Indian Telugu in the UK [ITU] - HG03968
- Bengali in Bangladesh [BEB] - HG04153, HG04155
- East Asia (EAS):

- Han Chinese in Beijing, China (HapMap) [CHB] - GM18552*, NA18570, NA18571
- Chinese Dai in Xishuangbanna, China [CDX] - HG00851*, HG01802, HG01804
- Kinh in Ho Chi Minh City, Vietnam [KHV] - HG02064, HG02067
- Japanese in Tokyo, Japan (HapMap) [JPT] - NA19068*
- Africa (AFR):

- Luhya in Webuye, Kenya (HapMap) [LWK] - GM19440*
- Gambian in Western Division, The Gambia [GWD] - HG02623*
- Esan from Nigeria [ESN] - HG03115*
- Europe (EUR):

- Toscani in Italia (Tuscans in Italy) (HapMap) [TSI] - GM20587*
- British from England and Scotland, UK [GBR] - HG00250*
- Finnish in Finland [FIN] - HG00353*
- America (AMR):

- Mexican Ancestry in Los Angeles, California, USA (HapMap) [MXL] - GM19789*
- Peruvian in Lima, Peru [PEL] - HG01971*

Asterisks next to sample IDs refer to samples processed at UCSF.

### Preprocessing 1000 Genomes Data

The 1000 Genomes data was separated into individual and chromosome specific VCFs using *vcftools^25^.* Further, the variants were filtered for biallelic SNPs, phased, filtered for PASS, and indels were removed. The experimentally phased data also had a very small fraction of unphased SNPs, which were removed by filtering with *vcftools.* The analysis was performed only for autosomes.

### Phasing Analysis

The alternate (ALT) allele frequencies of all the SNPs of interest were obtained from the 1000 Genomes data and converted to minor allele frequencies to be able to analyze switch error as a function of minor allele frequencies. The filtered SNPs from the experimental data were split into phase sets, based on phase set information available in the experimental VCF files. Switch error was calculated between the experimental and 1000 Genomes data for each phase set in each chromosome of each individual from the experimental dataset. Switch error is defined as *percentage of possible switches in haplotype orientation used to recover the correct phase in an individual^26^* or *proportion of heterozygous positions whose phase is wrongly inferred relative to the previous heterozygous position^27^. vcftools* returns the switch error as well as all positions of switches occurring along the chromosome.

#### Switch Error as a Function of Minor Allele Frequency

ALT allele frequencies were accessed for each of the switch positions from the data and were converted to minor allele frequencies. Distribution of switch positions as a function of minor allele frequency was plotted for each chromosome in each individual.

#### Switch Error as a Function of Inter SNP Distance

Positions of each SNP were accessed from the data. The number of intermediate switches were counted for all pair of SNPs, not only consecutive SNPs. If the number of switches between two SNPs were odd, a switch error was counted. This was used to calculate the distribution of switch errors as a function of inter-SNP distance.

### Imputation Analysis

The entire imputation analysis is performed for each chromosome for each individual.

#### Generate Recombination Map

*IMPUTE v2^13^* makes available recombination maps for each chromosome using the 1000 Genomes data for the GRCh37 assembly. A recombination map was obtained for each chromosome for GRCh38 by lifting over the GRCh37 maps using the *liftover* software. ~8k positions (0.2%) were removed from the lifted over recombination map because liftover resulted in them being in the incorrect order.

#### Generate Reference Panel

A reference haplotype panel was generated for all individuals from the 1000 Genomes data by subsetting it to the specific population of interest. 1000 Genomes data for the individuals which were experimentally sequenced was not included in the reference panel. *vcftools* was used to filter out the individuals of interest from the 1000 Genomes data. *bcftools* was used to convert the VCF data to haps-sample-legend format. An alternate approach was also used, where the entire 1000 Genomes data was used to generate a reference haplotype panel.

#### Generate Study Panel

A study panel was generated for the experimentally sequenced individuals selected. The study panel is assumed to be genotyped at positions corresponding to the Illumina **InfiniumOmni2.5-8** array. Array positions were lifted over from GRCh37 to GRCh38 using *liftover.* 1000 Genomes haplotypes (since 1000 Genomes data is prephased, the study panel is also in the form of haplotypes rather than genotypes) for those positions for those individuals were selected to create the study panel using *vcftools.* Filtered VCF files were converted to the haps-sample format using *bcftools.*

#### Run Imputation

Missing positions are imputed using *IMPUTE v2.* Imputation was performed in 5Mb windows. The genotype output by imputation was converted to VCF format using *bcftools.* VCFs produced over all windows were combined using *vcf-concat. IMPUTE v2* generally phases the typed genotyped sites in study panel. This is followed by imputation which is performed by assuming that haplotypes in the study panel that match the haplotypes in the reference panel at the typed sites also match in the untyped sites. *IMPUTE v2* then performs an iterative process performing multiple Monte-Carlo steps alternating phasing and imputation. For this analysis, however, as haplotypes from the 1000 Genomes project were directly used to generate the study panel, the phasing step was not performed.

#### Filter Positions

For one part of the analysis, i.e. estimating errors in the positions represented in the experimentally phased VCFs (henceforth called experimental SNPs), the positions from those VCFs were filtered from the imputed data using *vcftools.* Experimental genotypes from the experimental VCFs were obtained for each individual of interest using *vcftools.* SNPs with duplicate entries in either the imputed or experimental data were removed. Continent-specific allele frequencies were obtained for the experimental SNPs from the 1000 Genomes data using *vcftools*, to be able to analyze switch error as a function of Minor Allele Frequencies. For the other part of the analysis, i.e. estimating errors for all positions in the 1000 Genomes data, the allele fractions were similarly obtained for all of the SNPs.

#### Imputation Error

Imputation error was computed as fraction of genotypes being incorrectly identified. Imputation error was computed for both, the SNPs in the experimental data and all the SNPs in 1000 Genomes data. Error is computed as a function of minor allele frequency. The continent-specific minor allele frequencies were used for analyzing the imputation error.

For all analysis where error rate is computed as a function of the continent-specific minor allele frequency (genotyping error and imputation error; Figs. 1,2,7,8), the minor allele frequencies are binned as MAF=0.0%, 0.0% < MAF < 0.2%, 0.2% <= MAF <0.5%, 0.5% <= MAF < 1%, 1% <= MAF < 5%, MAF >= 5%. For the analysis where all 1000 Genomes minor allele frequencies are used (phasing error and imputation error comparing use of multiple reference panels; Figs. 3, 4, 9), the minor allele frequencies are binned into only five bins, i.e. there is no MAF=0.0% bin. Rest of the bins are the same as for the continent-specific MAF bins.

**Figure 1.**
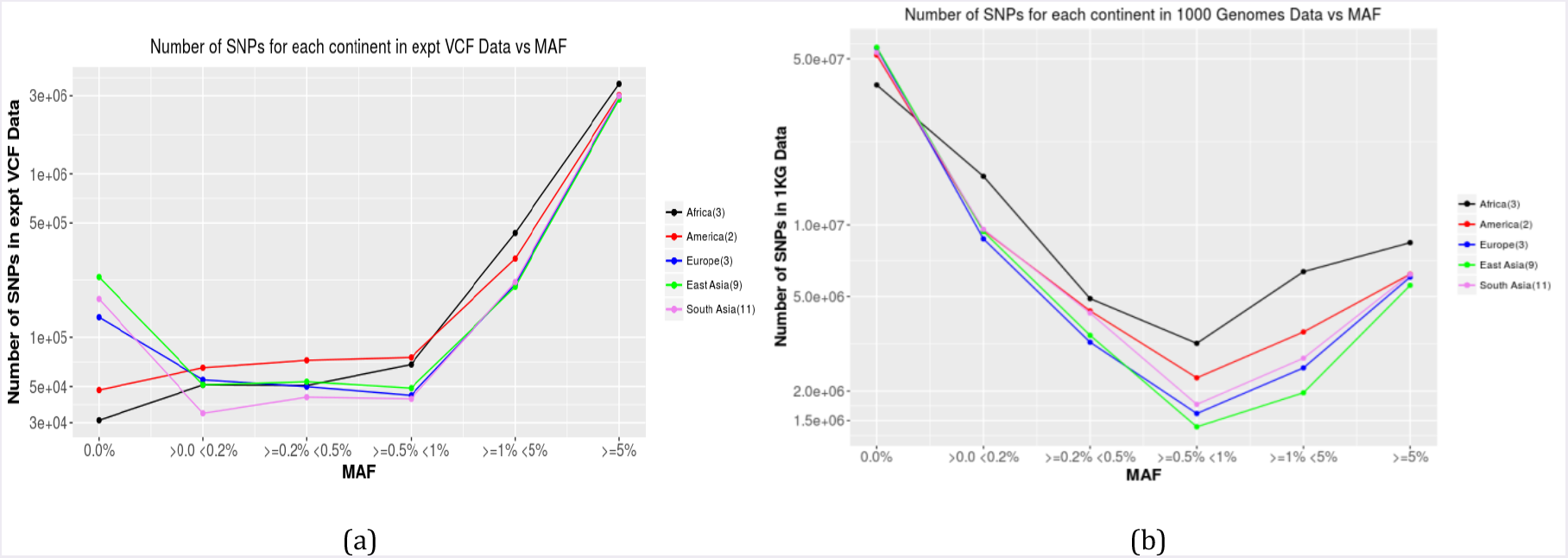
Distribution of SNPs as a function of continent-specific minor allele frequencies (a) only experimental SNPs (b) all 1000 Genomes SNPs

**Figure 2.**
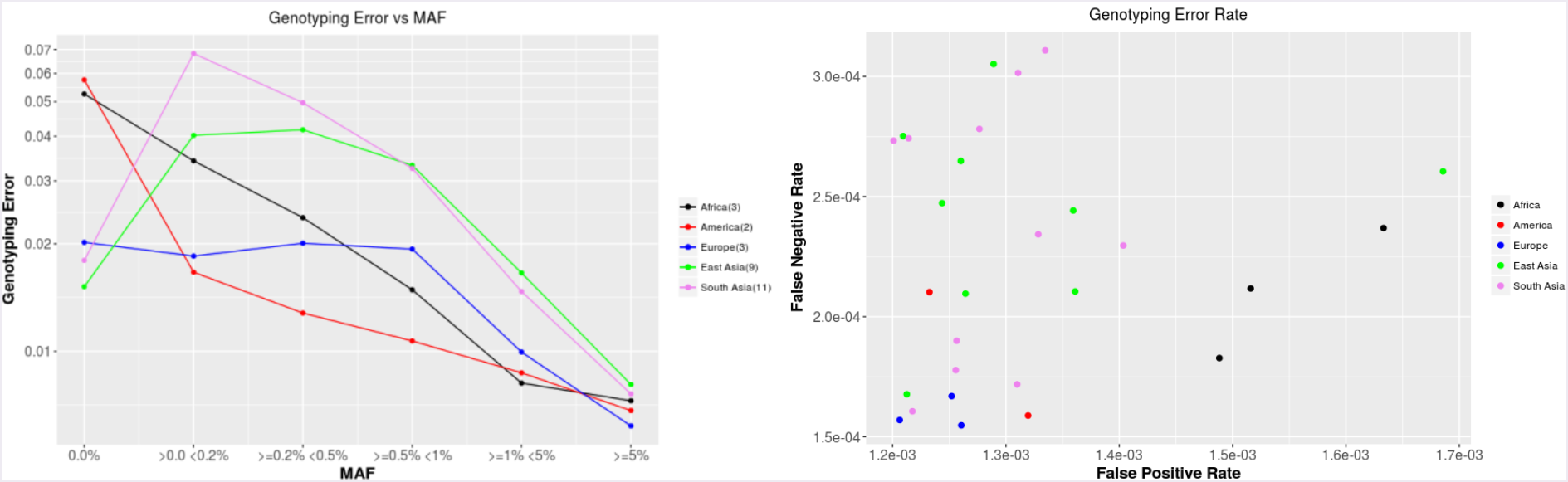
Genotyping error (a) in the experimental VCF positions as a function of continent-specific minor allele frequency averaged over all chromosomes over all individuals in each continent (b) false positive vs false negative rates (defined in text) for all 1000 Genomes SNPs

**Figure 3.**
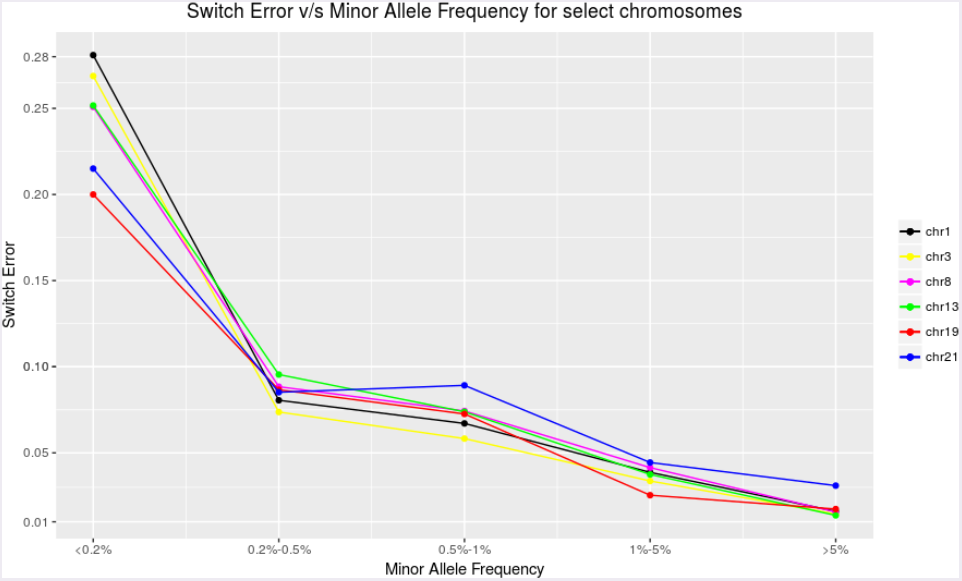
Switch error as a function of Minor Allele Frequencies for different individual chromosomes. Chromosome 21 shows higher switch error for large MAF values

**Figure 4.**
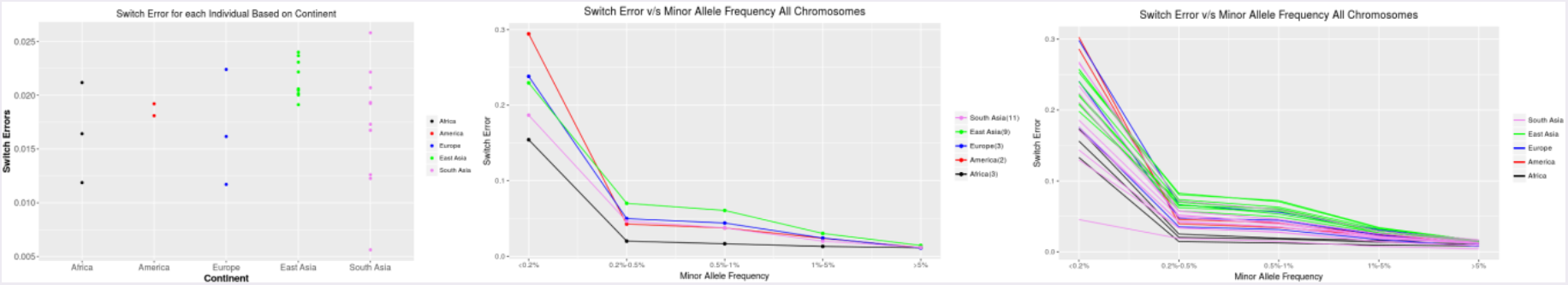
Switch error (a) Total switch error (number of switches in experimental SNPs/total number of experimental SNPs) for each individual (b) Switch error as a function of Minor Allele Frequencies averaged over all individuals in each continent. (c) Switch error as a function of Minor Allele Frequencies for all individuals colored by continent.

**Figure 8.**
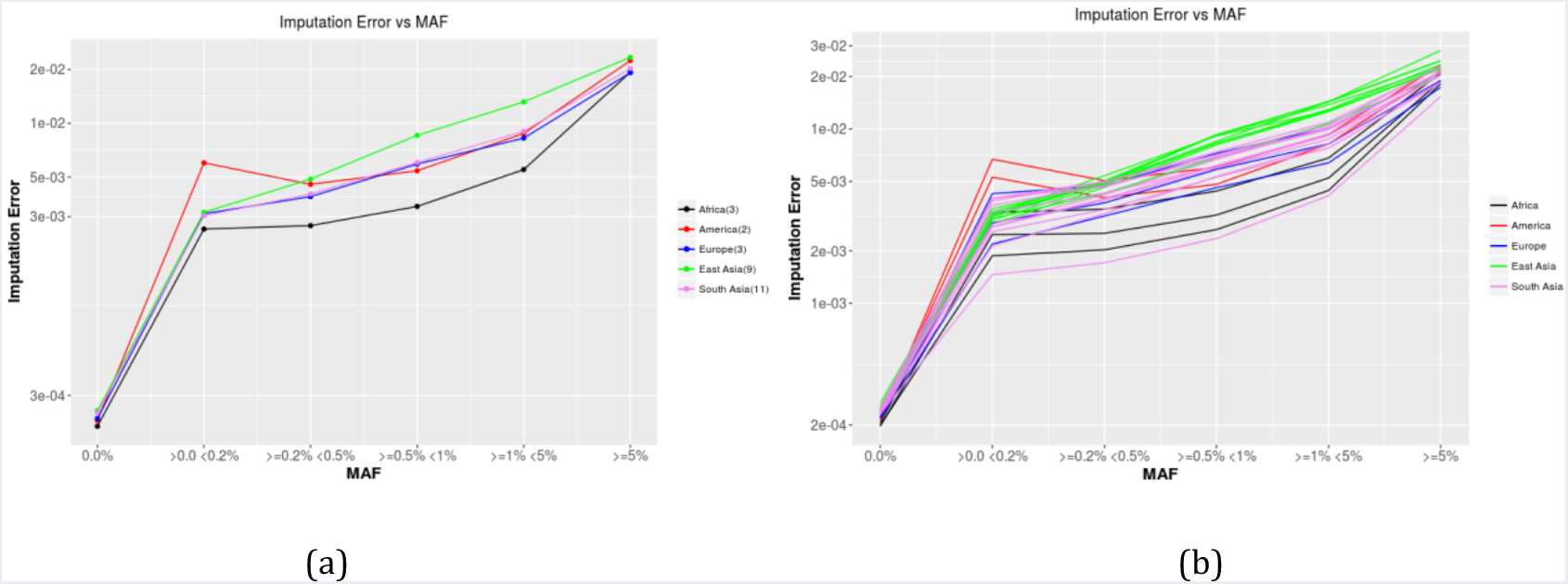
Imputation accuracy all 1KG SNPs (a) Imputation error in all the 1000 Genomes positions as a function of Minor Allele Frequencies averaged over individuals in each continent. (b) Imputation error in all the 1000 Genomes positions as a function of Minor Allele Frequencies for all individuals colored by continent.

**Figure 9.**
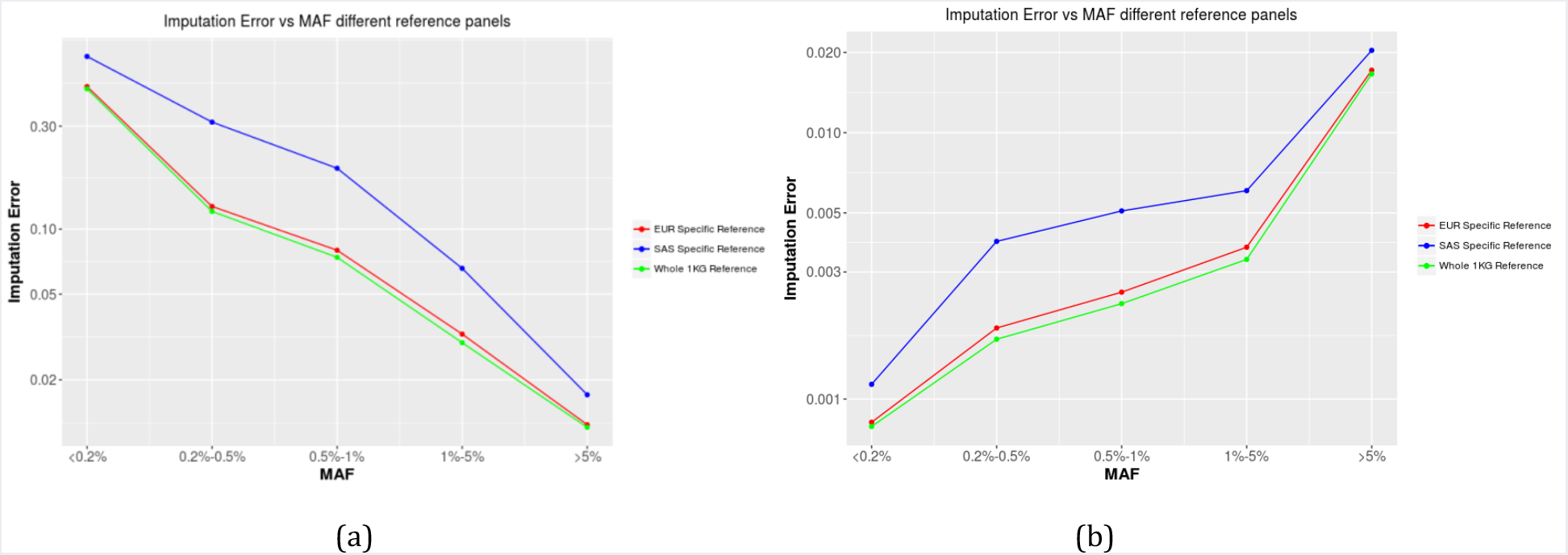
Imputation error as a function of Minor Allele Frequencies for European individuals comparing the European reference panel v/s the entire 1KG reference panel (a) experimental SNPs (b) All 1000 Genomes SNPs

### Experimental Methods

#### Samples processing

HMW Genomic DNA was extracted and converted into 10x sequencing libraries according to the 10X Genomics (Pleasanton, CA, USA) Chromium Genome User Guide and as published previously^28^. Briefly, GEMS were made with 1.25ng HMW template gDNA, Master-mix Genome Gel Beads and partitioning oil on the microfluidic Genome Chip. Isothermal incubation of the GEMs (for 3 h at 30°C; for 10 min at 65°C; stored at 4°C) produced barcoded fragments ranging from a few to several hundred base pairs. After dissolution of the Genome Gel Bead in the GEM Illumina Read 1 sequencing primer, 16bp 10x barcode and 6bp random primer are released. The GEMs were then broken and the pooled fractions were recovered. Silane and Solid Phase Reversible Immobilization (SPRI) beads were used to purify and size select the fragments for library preparation. Library prep was performed according to the manufacturer s instructions described in the Chromium Genome User Guide Rev C. Libraries were made using 10x Genomics adapters. The final libraries contain the P5 and P7 primers used in Illumina bridge amplification. The barcoded libraries were then quantified by qPCR (KAPA Biosystems Library Quantification Kit for Illumina platforms). Sequencing was done using Illumina HiSeq 4000 with 2×150 paired-end reads. Raw reads were processed, aligned to the reference genome, and had SNPs called and phased using 10X Genomics’ Long Ranger software (version 2.1.1 or 2.1.6) with the “wgs” pipeline with default settings.

## RESULTS

The 1000 Genomes project chromosome-specific VCFs for the GRCh38 assembly contain between 6.4M (chr1) to 1.1M (chr22) variants over all the 2504 individuals. After filtering for biallelic SNPs, phased, filtered for PASS, removing indels, we are left with 6.15M (chr1) to 1.05M (chr22) variants. The experimentally phased data from the 10X Genomics platform has different numbers of called variants for each sequenced individual. For chromosome 1, the number of called variants varies from 414K to 494K across the 28 individuals, while, for chromosome 22, the number of called SNPs varies from 104K to 120K. After performing a similar filtering for the experimental data, the number of biallelic PASS phased SNPs ranges between 298K and 357K for chromosome 1 and 64K and 75K for chromosome 22.

The SNPs from the experimentally phased VCFs (Fig. 1a), averaged over continent groups show that the vast majority of SNPs in this selection have high continent-specific MAF values (> 5%). Comparing across continents for the continent invariant SNPs, the African and American individuals have an order of magnitude less continent invariant SNPs than the European, East Asian and South Asian individuals. However, if we look at all the SNPs in the 1000 Genomes Data (filtered for biallelic PASS phased SNPs) as a function of continent-specific MAF, the distribution we observe has a very different trend. There is a significant over-representation of the very low continent-specific MAF SNPs (< 0.1%), ~ 5 * 10^7^, as compared to all the subsequent higher MAF SNPs, which all range < 1 * 10^7^.

These discrepancies between the numbers in the 1000 Genomes data and in the experimentally phased data, as well as the differing trends as a function of MAF occur because the 1000 Genomes data includes a SNP if even one individual in the 2504 individuals has a variant (heterozygous or homozygous-alternate) at that position while the experimental data includes a SNP only if that particular individual has a variant (heterozygous or homozygous-alternate) at that position. This results in a much larger number of overall SNPs being present in the 1000 Genomes data as compared to the experimental and also the majority of the 1000 Genomes SNPs having extremely low MAF, as those would occur only in one or a few individuals.

### Genotyping Error

Genotyping error is computed comparing the 1000 Genomes genotypes with the experimental genotypes. The experimental genotypes for all SNPs not present in the experimental VCF for each individual are assumed to be homozygous reference. Mismatched genotypes are counted as errors. Figure. 2a looks at the errors (fraction of genotypes which are incorrect) for the experimental VCF positions as a function of the continent-specific minor allele frequencies. There is higher error at the population invariant sites (MAF=0.0%) in the African and American populations than the European, East Asian and South Asian populations. This correlates with a lower total number of population invariant SNPs in those continents (Fig. 1a). For non-invariant SNPs, we observe, as expected, a decreasing error rate with increasing minor allele frequency, to a <1.5% error genotyping error rate for the SNPs with minor allele frequencies > 1%.

Comparing false positive (sites non-homozygous reference in 1000 Genomes data and homozygous reference in the experimental data) vs false negative (sites homozygous reference in 1000 genomes data and non-homozygous reference in the experimental data) error rates for all 1000 Genomes sites (Fig. 1b), we see that the genotyping for the European and American individuals is very accurate, with both low false positive and false negative rates. The East Asian and South Asian populations both have mostly low false positive rates, but show a wide range (factor of 2) of false negative rates, while showing only a ~15% variation in the false positive rates for most individuals. In contrast, the African individuals mostly have relatively low false negative rates, but have among the highest false positive rates. This indicates that the sequencing in the 1000 Genomes project has over called non-homozygous reference variants in African individuals compared to the rest, and over called SNPs as homozygous reference in some of the East and South Asian individuals.

### Phasing

Phasing errors are all analyzed for overall 1000 Genomes minor allele frequencies, not continent specific MAFs. Comparing the switch error across individual chromosomes (Fig. 3), we observe that the switch error ranges between 25 - 30% for the rare MAF (< 0.1%) SNPs, falling to < 5% for SNPs with MAFs 1 - 5%. The majority of SNPs, which fall in the MAF > 5% category, have an error < 2.5%. However, a comparatively higher switch error at larger MAF values (> 5%) is observed for chromosome 21. This plot (Fig. 3) shows only a subset of chromosomes a single individual (GM18552), but this trend is observed for all other chromosomes and individuals studied.

Figure. 4a shows the total switch error for each of the individuals. The total switch errors for all the individuals studied go up to ~ 2.5%. The switch errors for the East Asian individuals are grouped together, while those for the South Asian individuals show greater variability. This is in line with the general observation that South Asian populations have an overall greater heterogeneity than do East Asian populations [J. Wall, Unpublished data]

Analyzing the switch error as a function of minor allele frequency averaged over all chromosomes of all individuals of a population (Fig. 4b), we observe low switch error, < 5%, for low minor allele frequencies (MAF) (1 - 5%). For rare SNPs with MAF (0.2 - 1%), the switch error is ~ 5 - 10%. For extremely rare minor allele SNPs, i.e. MAF < 0.2%, the error is much higher, i.e. 15 - 35%. For all higher MAF values (> 5%), the error is < 2.5%. The average error rate for the individuals from the African populations is almost the same over the range of MAF values > 0.1%.

As observed in Figure. 4c, the differences in the error rates between individuals decrease with increasing minor allele frequency. Individuals from South Asia show a larger variation in error as a function of MAF as compared to individuals from East Asia. The individuals from the African populations have the lowest switch error over the range of MAF values. Individual NA20900, an individual from the Gujarati Indians in Houston (GIH) population has the lowest switch error as a function of minor allele frequency for the low MAF SNPs.

We also analyzed phasing error as a function of the distances between SNPs (Fig. 5). The phasing error increases as a function of the inter-SNP distance, i.e. SNPs which are further apart are more likely to be out of phase with each other. The within population trends are the same as for switch error vs MAF, where the individuals from South Asia show a larger spread as compared to the individuals from East Asia. Individual NA20900 shows the lowest error rate, same as for the comparison of error vs MAF (Fig. 4c).

**Figure 5.**
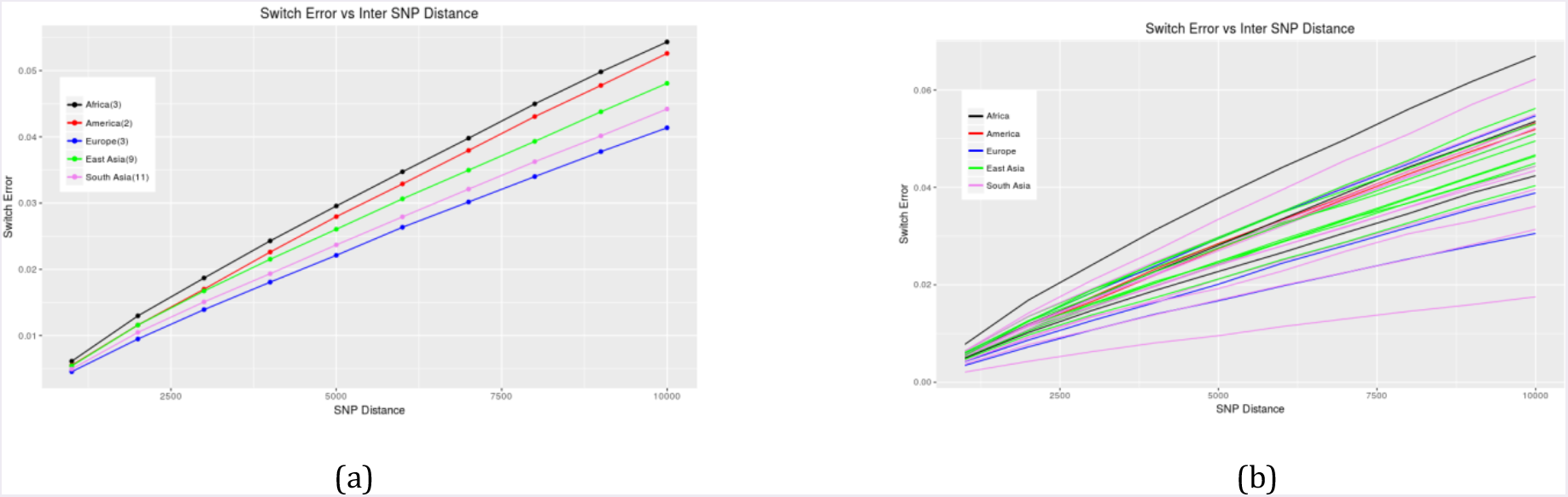
Switch error as a function of inter-SNP distance (a) Switch error as a function of inter-SNP distances averaged over individuals in each continent. (b) Switch error as a function of inter-SNP distances for all individuals colored by continent.

Comparing the switch error as a function of MAF vs. the switch error as a function of inter-SNP distance, we see that the individuals from the African populations show distinctly opposite trends. For low MAF SNPs, the error is the lowest averaging over the African individuals, while across the range of inter-SNP distances, the average over the African individuals was the highest error. The reason this occurs can be understood from the fact that there are a higher number of low MAF SNPs in the African individuals in the experimental data (Fig. 1a), as well as an overall higher number of SNPs in those individuals, leading to a higher SNP density for these individuals. In addition, there is less linkage disequilibrium (LD) in the individuals from the African populations, which would make it harder to phase them accurately^29-30^. Hence, pairs of SNPs are more likely to be out of phase with each other, leading to higher switch error as a function of inter-SNP distance.

**Imputation**

Imputation error is computed as the fraction of SNPs with incorrectly imputed genotypes. However, depending on the subset of SNPs under consideration, the error can be computed in two different ways, (1) fraction of experimental SNPs incorrectly imputed and (2) fraction of all 1KG SNPs incorrectly imputed. In the case of the second definition of error, the experimental calls for all the positions not in the experimental VCFs are set to homozygous-reference.

Figure. 6a shows the total imputation error in the experimental SNPs while Figure. 6b shows the total imputation error in the 1KG SNPs for each of the individuals. The total imputation errors in the experimental SNPs for all the individuals studied go up to ~ 4%. For this subset of SNPs, the two American individuals have the among the highest imputation errors. The imputation errors for the East Asian individuals are grouped together, while those for the South Asian individuals show greater variability. This agrees with our observations for the switch error (Fig. 4a). In the 1KG SNPs, on the other hand, since we are looking at a much larger set of SNPs, most of which are homozygous-reference in any given individual, we see a much smaller error <~ 1%.

**Figure 6.**
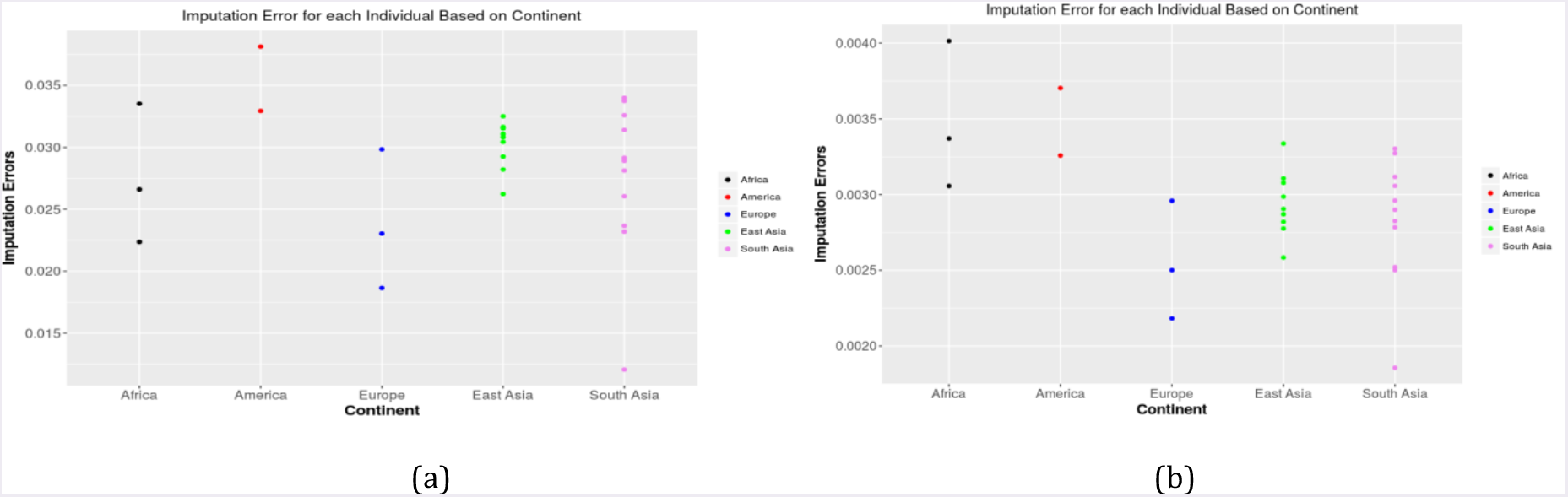
Total imputation error (a) Total imputation error in experimental SNPs (number of incorrect genotypes in all experimental SNPs/total number of experimental SNPs) for each individual (b) Total imputation error in all 1KG SNPs (number of incorrect genotypes in all 1KG SNPs/total number of 1KG SNPs) for each individual

**Figure 7.**
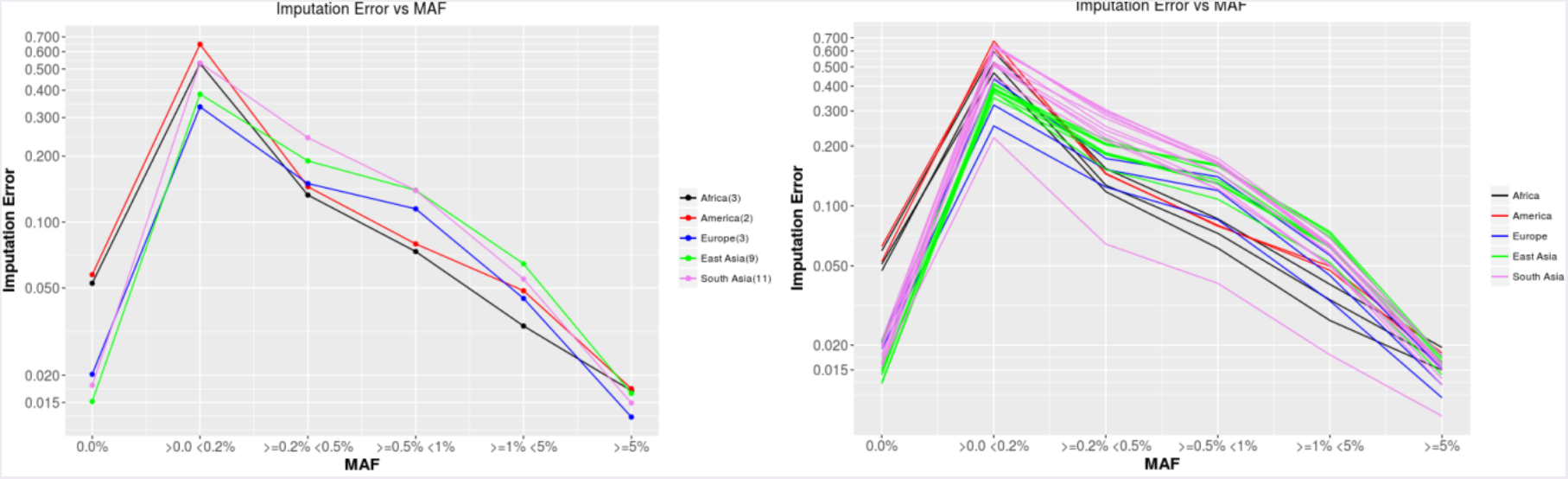
Imputation accuracy experimental VCF positions (a) Imputation error in the experimental SNPs as a function of Minor Allele Frequencies averaged over individuals in each continent. (b) Imputation error in the experimental SNPs as a function of Minor Allele Frequencies for all individuals colored by continent.

### Imputation error in experimental SNPs

Figure. 7a shows a wide range of error rates as function of the continent-specific minor allele frequency. The continent invariant positions (MAF=0.0%) are imputed almost as accurately as the high MAF (>5% in 3 populations, and >1% in two populations) SNPs. In these positions, we make the same observation as we did for the original genotyping in the 1000 genomes reference data (Fig. 2a), i.e. the errors in the European, East Asian and South Asian individuals for these continent invariant positions are lower than those for the American and African individuals. For the very rare SNPs, i.e. MAF < 0.2%, the error is as high as ~ 60%. These extremely high error rates are only observed in the American individuals and a few of the South Asian individuals. For the rest of the individuals, the error rates are < 50%. In the mid-range of MAF values, i.e. 0.2% to 1%, the errors range between 10 - 20%. The SNPs with higher MAF values are fairly accurate, with errors < 2% for common SNPs (MAF > 5%). This can also be seen looking at all the individuals separately (Fig. 7b). The South Asian (Gujarati in Houston, Texas) individual NA20900 still shows the lowest error rate as a function of MAF for imputation, just as it does for the switch error (Fig. 4c).

### Imputation error in all 1KG SNPs

Computing the error using all the 1KG SNPs, we see a different trend for the errors as a function of minor allele frequency (Figs. 8a, 8b). The invariant sites have very low errors ~10-^4^. For the variant sites, the errors increase as a function of minor allele frequency, as opposed to decreasing as they do in the experimental only SNPs. The reason this happens is that contrasting the number of experimental SNPs (Fig. 1a) with the numbers of all 1KG SNPs(Fig. 1b), while the number of low MAF SNPs is 1-2 orders of magnitude less than the number of SNPs with MAF >

5% in the experimental data, the number of very low MAF SNPs is 2-10 times greater than the number of SNPs with MAF > 5% in the whole 1000 Genomes data. The vast majority of the very low MAF SNPs in the whole 1000 Genomes data are homozygous-reference, since those SNPs show variation in only one or very few 1000 Genomes individuals. Hence, imputation predictions get most of those positions correct in most of the individuals. As a result, the fraction of those very rare SNPs which are predicted incorrectly is much lower when considering all the 1000 Genomes SNPs as compared to only considering the experimental SNPs, where most of the SNPs are high MAF SNPs.

Consistent with the observations for the experimental only SNPs, at very rare SNPs (MAF < 0.2%), the American individuals still have the highest error rate. The individuals from the South Asian populations still show a greater spread than those from the East Asian populations. Individual NA20900 still shows the lowest error rate as with previous observations.

### Comparison of reference panels

Here, we compare the imputation errors resulting from using different reference panels for imputation. A continent-specific reference panel for the individual of interest, a reference panel which includes all of the 1000 Genomes individuals, and a continent-specific reference panel for a different continent from the one from which the individuals are, are chosen. The minor allele frequencies used here are for all the overall 1000 Genomes minor allele frequencies, instead of a continent-specific minor allele frequency, since we want to understand the impact of the choice of reference panel, and continent-specific MAFs would not align with the whole reference or the reference from another continent. In this case, we look at the imputation error in the 3 European individuals when imputation is carried out with the European reference, the South Asian reference, and the whole 1000 Genomes reference.

The observed result for experimental only SNPs (Fig. 8a) when comparing reference panels for the European individuals is very similar when looking at all 1000 Genomes SNPs (Fig. 8b). The imputation accuracy when using the entire 1000 Genomes data as a reference panel gives a slightly better accuracy than using just a European specific reference panel. The error while using an incorrect reference panel, however, is up to a factor of 2 greater than the error when using the appropriate reference, or when using the whole 1000 Genomes reference panel. The trend of error as a function of MAF is, again, the opposite of what was observed when looking at only the experimental SNPs.

## DISCUSSION

The 1000 Genomes Project data have been widely used as a reference for estimating continent-specific allele frequencies, and as a reference panel for phasing and imputation studies. Since the project’ s design involved low-coverage (~7X) sequencing for most of the samples, it was unknown *a priori* how accurate the 1KGP’ s genotype and haplotype calls were, especially for rare variants. This accuracy obviously directly impacts the usefulness of the 1KGP data. With the advent of inexpensive, commercial platforms for experimentally phasing whole genomes, it is possible to directly quantify the genotype and haplotype error rates of the 1KGP data.

Our comparison of 28 experimentally phased genomes with the 1KGP data found that the latter is highly accurate for common and low-frequency variants (i.e., MAF ≥ 0.01). As expected, accuracy declined with decreasing MAF, with rare variants (MAF < 0.01) not reliably imputed onto haplotypes. Surprisingly though, the genotype calls were reasonably accurate even for rare variants. This observation may not generalize to other low-coverage sequencing studies due to the complicated and labor-intensive protocol used for variant calling in the 1KGP. We conclude that the 1KGP data is best used as a reference panel for imputing variants with MAF ≥ 0.01 into populations closely related to the 1KGP groups, and is probably of limited utility for imputation in rare variant association studies. Larger subsequent imputation panels, such as the one generated by the Haplotype Reference Consortium (HRC)^31^, are likely much more useful for imputing rare variants, at least in well-studied European populations. However, even this large reference panel may be of limited usefulness for imputation into other human groups. While our results suggest that using a region-specific reference panel (for the correct region) for imputation is only slightly worse than using a worldwide panel, the choice of an incorrect regional panel makes the imputation considerably worse. So, large European-based haplotype reference panels will be of limited utility for imputing variants into East Asian, South Asian, or African-American genomes, while imputation studies involving understudied groups such as Middle Easterners, Melanesians or Khoisan are likely to have error rates substantially higher than what was observed in our study. This is a consequence of the fact that most rare variants are region-specific; imputation only works when the variant being imputed shows up often enough in the reference panel. In summary, while the 1KGP and HRC provide valuable genomic resources that can augment the power of GWAS in groups with European ancestry, additional large-scale genome sequencing of diverse human populations will be necessary to obtain comparable benefits of imputation in genetic association studies of non-European groups.

Finally, we note that the absolute error rate varied by an order of magnitude, depending on the specific definitions of error that were used. This highlights the importance of definitional clarity in studies that evaluate the accuracy of genomic resources.

## Acknowledgments

JDW was supported in part by NIH grant R01 GM115433. SB was supported by Genentech research grant CA0095684.

